# Putting value in context: A role for context memory in decisions for reward

**DOI:** 10.1101/033662

**Authors:** Aaron M. Bornstein, Kenneth A. Norman

## Abstract

How does experience inform decisions? In episodic sampling, decisions are guided by a few episodic memories of past choices. This process can yield choice patterns similar to model-free Reinforcement Learning (RL); however, samples can vary from trial to trial, causing decisions to vary. Here, we show that context retrieved during episodic sampling can cause choice behavior to deviate sharply from the predictions of RL. Specifically, we show that, when a given memory is sampled, choices (in the present) are influenced by the properties of other decisions made in the same context as the sampled event. This effect is mediated by fMRI measures of context retrieval on each trial, suggesting a mechanism whereby cues trigger retrieval of context, which then triggers retrieval of other decisions from that context. This result establishes a new avenue by which experience can guide choice, and as such has broad implications for the study of decisions.

## Introduction

How do we learn from our past decisions? According to the dominant model-free reinforcement learning (RL) theory of choice, actions are selected based on expected values that are computed as running averages of experienced rewards. This average is updated incrementally as new rewards are incorporated, resulting in a steadily decaying influence of past experiences [1]. We have previously shown that this pattern of dependence on experience can also result from an active deliberation process that draws, in a recency-weighted fashion, on episodic memories of relevant past choices as samples of possible outcome values [2]. For instance, we might evaluate a restaurant by recalling recent dining experiences at similar establishments.

Both approaches assume that the influence of a past event on choice is a simple function of how long ago that event was experienced or remembered. Where they differ is in how that influence arises. In RL, the contribution of a given past trial to reward estimates on a given choice is a fixed, decreasing function of its age. In episodic sampling this influence is dynamic, which can cause its choices to diverge from RL. Because the process draws only a small number of samples, a given episode (even sometimes one from the far past) will, when recalled, have a large contribution to the estimated value for that decision. At the same time, recent episodes could be overlooked, and thus have no influence on the current choice. This distinction can be obscured when looking at average choice behavior in the sort of repeated decision task usually employed in the laboratory [3]. The difference between the predictions of these two models is more pronounced when incidental reminders of past choices are introduced to the decision-making task [2]. These reminders cue the past trial episode, recalling the action taken and reward received, and thus affect the next decision in a way not captured by standard RL.

However, episodic memories consist of more than just the simple association between action and outcome. They also carry rich information about the temporal, spatial, and visual *context* of an experience [4,5]. When context is reinstated, it affects what we remember next: after recalling one event, we are more likely to subsequently recall events that share context with the first [6]. For instance, when recalling one restaurant, we might also recall the street it was on, which could lead to recalling another restaurant from the same street.

In this way, retrieval of contextual information could impact decisions made by episodic sampling. Specifically, context could induce a form of autocorrelation in sampling: The first sample also brings to mind the context from which subsequent samples are likely to be drawn. These following samples would also have an impact on decisions. The average influence on choice of a past episode would therefore be a function both of its age and the probability that other, contextually-related episodes would bring it to mind. This extra influence of context (if present) would constitute a radical departure from incremental RL, which has no means of accounting for this influence.

To probe whether, and by what mechanism, context biases episodic sampling, we designed an experiment to isolate the effects of retrieved context on decision-making, distinct from the effect of the initial sampled trial. Participants performed a three-option choice task in which trials took place across seven visually-distinct contexts, described as “rooms” of a virtual casino, each distinguished by a context-specific image of an outdoor scene. After making each choice, participants were shown both the reward they earned and a trial-unique object picture. Some of these objects were later presented during recognition memory probes that were interleaved with the choice trials. Importantly, we designed the experiment such that the rewarded choice associated with a probed object was different from the choice that was most frequently rewarded in the room (context) where the probed object was originally presented. This procedure allowed us to disentangle the influence of the reminded trial episode [2] from that of the context.

We hypothesized that, if recall of the reminded trial also triggered the reinstatement of context, we would observe that choices are influenced by the action rewarded in the context as a whole, not just the action rewarded on the reminded trial itself. This hypothesis further implies that the effects of retrieved context on choice will only be evident on trials where the object’s context (room) is retrieved. To test this prediction, we carried out an fMRI experiment and employed multivariate pattern analysis (MVPA) to covertly measure neural evidence for context reinstatement on each trial. Pattern classifiers (trained to recognize scene-related activity) output a trial-by-trial measure of how likely it was that participants were recalling a past context. We used this neural reinstatement index as a mediating variable to predict the effect of context on choices.

Experiment 1 provided a behavioral test of influence of context. Experiment 2 provided both a behavioral test and a neural test (using fMRI) of the predictions outlined above. Taken together, these experiments reveal new aspects of the computational and neural mechanisms by which individual episodes of past experience are brought to bear on decisions for reward, and introduce a novel signature of decisions guided by episodic memory.

## Results

In Experiment 1, 20 participants performed the task (Figure 1) and their behavior was analyzed for evidence of context’s influence on decisions. In Experiment 2, 32 additional participants performed the task while being scanned in fMRI, which allowed us to examine a neural mechanism that gives rise to – and predicts the degree of – the influence of context on decisions.

**Figure 1:**
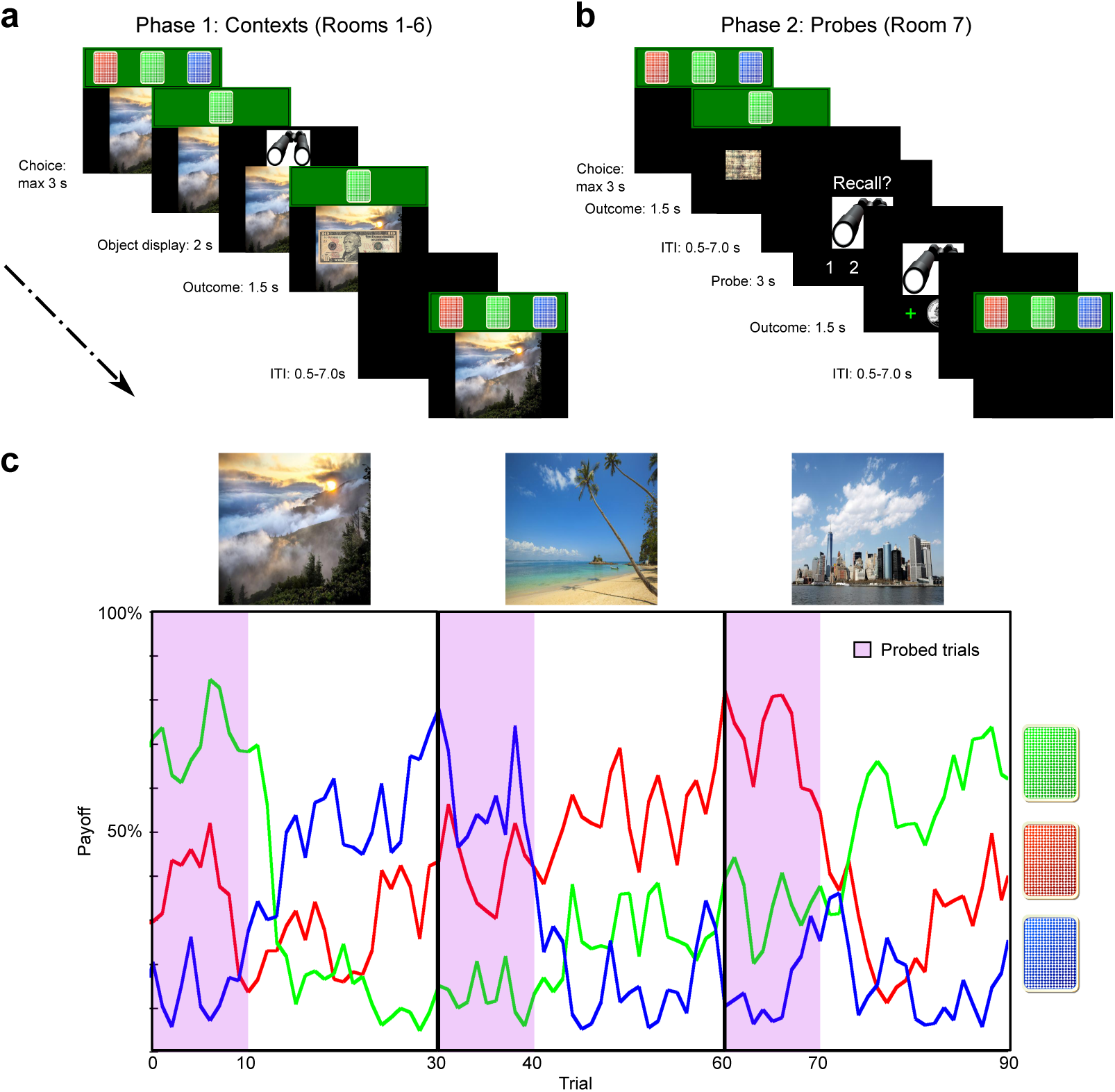
Task design. Participants (Experiment 1: 20; Experiment 2: 32) performed 300 trials of a three-option sequential choice task. **a**. **Context learning.** On each trial, participants chose between three card decks, each with a different, unsignaled, probability of paying out a $10 reward, as opposed to $0. After a deck was chosen, the top card was turned over, revealing a trial-unique object photograph. The first 180 choices took place in six “rooms” of a virtual casino. Rooms were distinguished by background photographs of outdoor scenes. **b. Memory probes.** The final 120 choices took place in a seventh room, which did not have a scene photograph in the background. Choices in this room were also rewarded, but did not result in object pictures. Interspersed among choices trials were 60 memory probe trials, which whether, and with what confidence, subjects remembered a given object picture. **c. Example payoff timeseries**. The probability that each deck would pay out $10 changed on each trial according to a Gaussian random walk centered around one of three distinct values: 60%, 30%, and 10%. Purple bands denote the first 10 trials in each new context room; after the 10th trial, the center values were shuffled across decks, such that the previous highest-paying deck was no longer the best option. Critically, images selected for memory probes were drawn only from these first 10 trials in each room, distinguishing the reward values of probed trials from those of the rest of the context.

The following three phases were common to both experiments. In *Phase 1*, participants performed 300 trials of a three-option rewarded choice task (Figure 1a,b). Choices returned either $10 or $0 with varying probability (Figure 1c; Equation 1). The first 180 trials took place across six “rooms” (contexts), distinguished by the presence of one of six scene images in the background (Figure 1a). In *Phase 2*, participants visited a seventh room, where no scene images were visible, and made 120 further choices; 60 recognition memory probes were interspersed between these choices at pseudorandom intervals (Figure 1b). On recognition probes, participants were asked whether, and with what confidence, they recognized the presented picture. Participants were rewarded with $0.25 for correct responses, and penalized by the same amount for incorrect responses. The average lag between initially viewing a picture and being tested on recognition of that picture was over 170 trials (Expt 1: Average lag 173.40 trials SEM 1.70; Expt 2: 175.88 SEM 0.64). In *Phase 3*, participants were given a source recognition test that assessed whether they could remember the room (context) in which the probed objects were encountered during Phase 1. Experiment 2 also had a fourth phase, an fMRI visual category localizer task used to train the pattern classifiers.

### Memory tasks

Participants performed well on the recognition memory probes in Phase 2 (mean d’, Experiment 1: 2.51 SEM 0.29, Experiment 2: 2.43 SEM 0.20). Trials with incorrect answers on the recognition memory probe were rare. Trials with incorrect or low-confidence answers were excluded from further analysis because they were not of interest for our hypothesis; our goal was to evaluate the effect of *successful* reminders on subsequent decisions, and low-confidence and/or incorrect responses indicated that the reminders were unsuccessful. Performance on the Phase 3 source recognition task was also well above chance (Experiment 1 presented all six options, so chance level was 16.67%: actual performance mean 45.80% SEM 5.38% correct; Experiment 2 subselected three options to fit the MRI button box, so chance level was 33.33%: actual performance mean 68.12% SEM 2.38% correct).

### Experiment 1

Our primary measurement of interest was performance on choice trials after the recognition memory probes. By our hypothesis, these trials should show a significant influence of rewards received on trials across the reminded context (i.e., the room in which the reminded trial occurred during Phase 1; Figure S1 and Figure S2).

We ran a multiple regression to model the effect on choice behavior of the recently received rewards, the identity of the recently chosen options, the value of the reward received on the probed trial, and the context reward. This analysis identified significant and separable effects of each of the three sources of reward information (Figure 2a); in particular, we found that memory influenced choice in two distinct ways. First, replicating our previous results [2], we found that the reward content of trials evoked by memory probes influenced the option selected by participants on the ensuing choice trial. If the probed trial was not rewarded, participants were less likely to choose as they had on that probed trial. The reward received on the probed trial was a significant predictor of choice (*t*(19)=2.2043, *P*=0.04), with a mean regression weight of comparable magnitude to that of rewards directly received three trials earlier.

**Figure 2:**
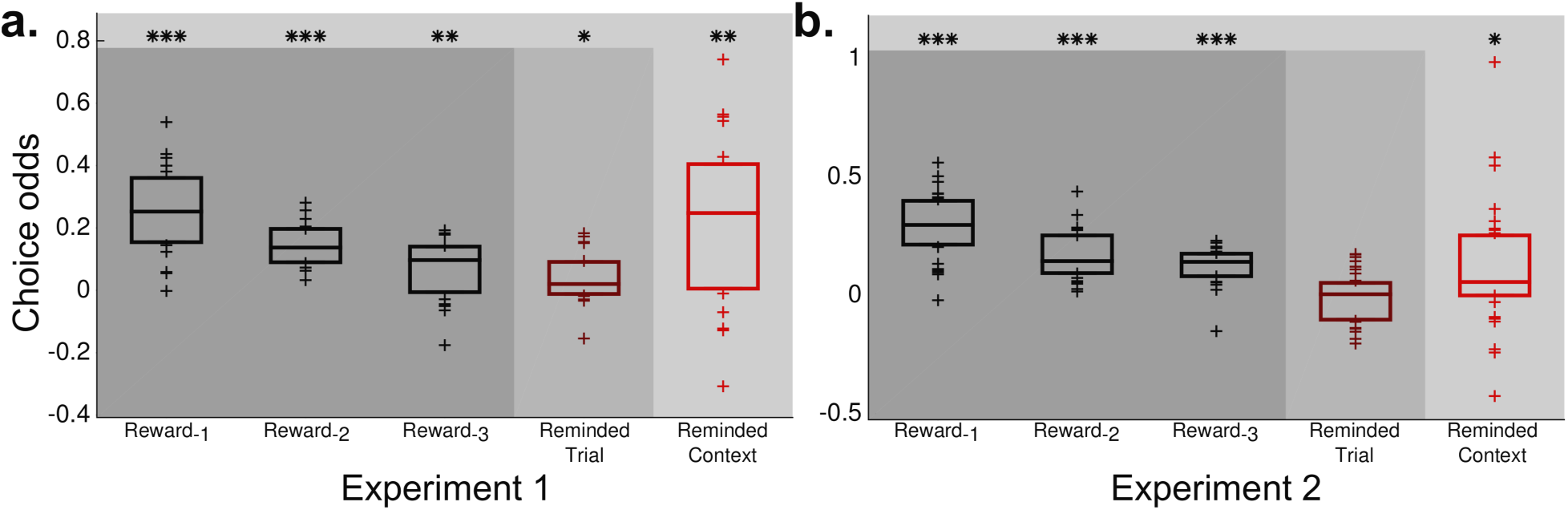
Context reward influences choices following a probe. Multiple linear regression of three sources of reward information on choices following a probe. The increase in choice weight for a given deck resulting from each reward term are plotted here as the median (+/− one quartile) across participants, separately for each experiment (**a** for Experiment 1, **b** for Experiment 2). Columns 1-3 show the effect of recent rewards, which reflect an exponentially decaying influence of recent experience of the sort that could arise from either incremental model-free reinforcement learning or recency-weighted sampling models. These effects were significant in both experiments (Expt. 1: R_-1_: *t*(19)=7.8678, *P*<0.0001; R_-2_: *t*(19)=9.5878, *P*<0.0001; R_-3_: *t*(19)=3.0066, *P*=0.0073; Expt. 2: R_-1_: *t*(31)=11.8339, *P*<0.0001, R_-2_: *t*(31)=9.2157, *P*<0.0001; R_-3_: *t*(31)=9.6940, *P*<0.0001). Column 4 shows the effect of reward received on the probed trial. This effect was significant in Experiment 1 (*t*(19)=2.2043, *P*=0.04), but not Experiment 2 (*t*(31)=−0.4878, *P*=0.6291). Column five shows the effect of rewards received in the context of the probed trial. This effect was significant in both experiments (Expt. 1: *t*(19)=3.5488, *P*=0.0021; Expt. 2: *t*(31)=2.4457, *P*=0.019). (* *P*<0.05, ** *P* < 0.01, *** *P* < 0.001)

Expanding beyond the previous results, we also found an effect of the rewards received for a given deck across other trials within the context of the probed trial (hereafter, the *context reward*; Equation 2). On choice trials following a memory probe, participants were more likely to choose a deck the greater were its proportion of trials being rewarded across the reminded context. This context reward was also a significant predictor of choices after a probe (*t*(19)=3.55, *P*=0.0021), with a mean regression weight of comparable magnitude to that of the reward received for direct experience just one or two trials previous. The effect of the reminded context was greater than the effect of the reminded trial (*t*(19)=2.7262, *P*=0.0134).

### Experiment 2

We then repeated the behavioral task from the first experiment with a new group of 32 participants. In this version, participants underwent fMRI scanning to allow us to identify brain activity predictive of the context reward effect. The results support the hypothesis that evoked context has a separate and strong influence on choice (Figure 2b).

Critically, as in the first experiment, the reward from the reminded context again had a significant effect on subsequent choice (*t*(31)=2.4457, *P*=0.0190). As in Experiment 2, the effect of reminded context was again greater than the effect of the reminded trial (*t*(31)=2.2613, P=0.0309). However, diverging from the results observed in Experiment 1 and the preceding study [2], the reward received on the probed trial did not have a significant effect on subsequent choice (*t*(31)=−0.4878, *P*=0.6291). Post-hoc simulations confirmed that both the relative prominence of the reminded context effect over the reminded trial effect and the sparing of the context effect when the single trial effect is not statistically reliable are consistent with our context-aware sampling mechanism. The intuition here is that – while the probe initially triggers sampling of the reminded trial – subsequent samples taken from the reminded context will outweigh the effects of sampling the reminded trial (Figures S3 and S4).

#### Regression results are incompatible with incremental learning models

For the analyses reported above, we designed our context reward regressor to capture the average effect of sampling memories from the probed context: For a given context, each episode in that context in which a given card deck was chosen was treated as evidence for or against choosing that card deck (depending or whether the choice was rewarded). This way of computing context reward is qualitatively different from that which would be predicted by an incremental learning algorithm such as model-free temporal-difference learning. This is because those incremental algorithms would more heavily weight later experiences, and those with higher reward prediction errors. However, in our formulation, every trial from the probed context has equal weight.

To explore whether the observed context reward effect could be explained by incrementally-learned action values, as in model-free reinforcement learning, we first fit RL models to choices in each of the six context rooms. Each model learned cached action values for the three card decks; two of these models reset those values when context changed–in one model, the context shifts/value resets at the time that the room changed, in the other model, values were reset at a variable trial number after the start of each room (to account for the possibility that context boundaries were inferred at the time the payoffs changed).

The model that reset action values when the room changed was the best fit to behavior. By BIC versus the second-best model, in Experiment 1, the room-reset model was superior for 18/20 subjects, mean difference in BIC: 5.6981; In Experiment 2, 26/32 subjects, 4.6406. The fit parameters for the room-reset model – learning rate α, softmax temperature β, choice stickiness β_p_ – were: Expt. 1 α mean 0.4802 SEM 0.0617, β mean 0.2333 SEM 0.2829, β_p_ mean 0.4348 SEM 0.1392; Expt. 2 α mean 0.5738 SEM 0.0375, β mean 0.4623 SEM 0.0313, β_p_ mean 0.2231 SEM 0.0738. These values were consistent across the two experiments (by unpaired, two-sample *t*-test: α: *t*(50)=1.3775, *P*=0.1745; β: *t*(50)=−1.0137, *P*=0.3156, β_p_: *t*(50)=1.4694, *P*=0.1480).

We took the final values computed by this model for each card deck in each context and used them as context reward regressors in a regression analysis following that of Figure 2. The mean correlation between this new regressor and the corresponding original regressor was *R*=0.0564 (SEM 0.0645, *t*(19)=0.8749, *P*=0.3926) in Expt. 1 and *R*=0.1714 (SEM 0.0313, *t*(31)=5.4768, *P*=5.4728e−06) in Expt. 2. In each experiment the effect of this RL-derived context reward (*β*_RLCR_) was significant or trending, but, critically, negative (Expt. 1, *β*_RLCR_=−0.0125, SEM 0.0058, *t*(19)=2.1484, *P*=0.0448 across subjects; Expt. 2, *β*_RLCR_=−0.0091, SEM 0.0050, *t*(31)=−1.8156, *P*=0.0791). When both the original context reward (β_CR_) and this new RL version were run alongside each other in a simultaneous regression, the weights to the original were unchanged compared to when run alone (Expt. 1: in the simultaneous regression, β_CR_=0.2335, SEM 0.0643; the difference between this and the original was not significant, *t*(19)=−0.1710, *P*=0.8660. Expt. 2: in the simultaneous regression, β_CR_=0.1306, SEM 0.0427; the difference between this and the original was not significant, *t*(31)=−0.4515, *P*=0.6548). Therefore, incrementally-learned action values cannot explain our observed results.

#### fMRI analysis

Previous work in our lab has shown that classifiers trained to identify fMRI correlates of scene processing can be used to track mental reinstatement of contexts in which scenes had (previously) been presented; furthermore, these neural measures of context reinstatement predict memory behavior [7]. We therefore used this same strategy in our study, “tagging” some contexts with scene pictures and then using scene evidence as a covert neural measure of context reinstatement. As both the probe image and the seventh “room” in which probes were presented are devoid of scene images, we interpret evidence of scene processing on probe trials as indicative of memory reinstatements, in particular of the scenes presented during the first six rooms of the experiment. On this basis we hypothesized that, as scene evidence increased, so too would the effect on decisions of context reward.

We first identified regions of bilateral posterior parahippocampal cortex that were preferentially activated by the processing of scene images, using a post-task localizer scan where participants viewed scene images (and other kinds of images) that were not in the experiment (Figure 3a). We then selected post-probe timepoints on which to perform our analysis. We selected as timepoints of interest those volumes following the presentation of a memory probe that reliably showed elevated classifier evidence for scenes (Figure 3b). Based on this measure, timepoints four through six – representing the period from approximately eight to approximately twelve seconds after the onset of the probe image – were selected as our timepoints of interest.

**Figure 3:**
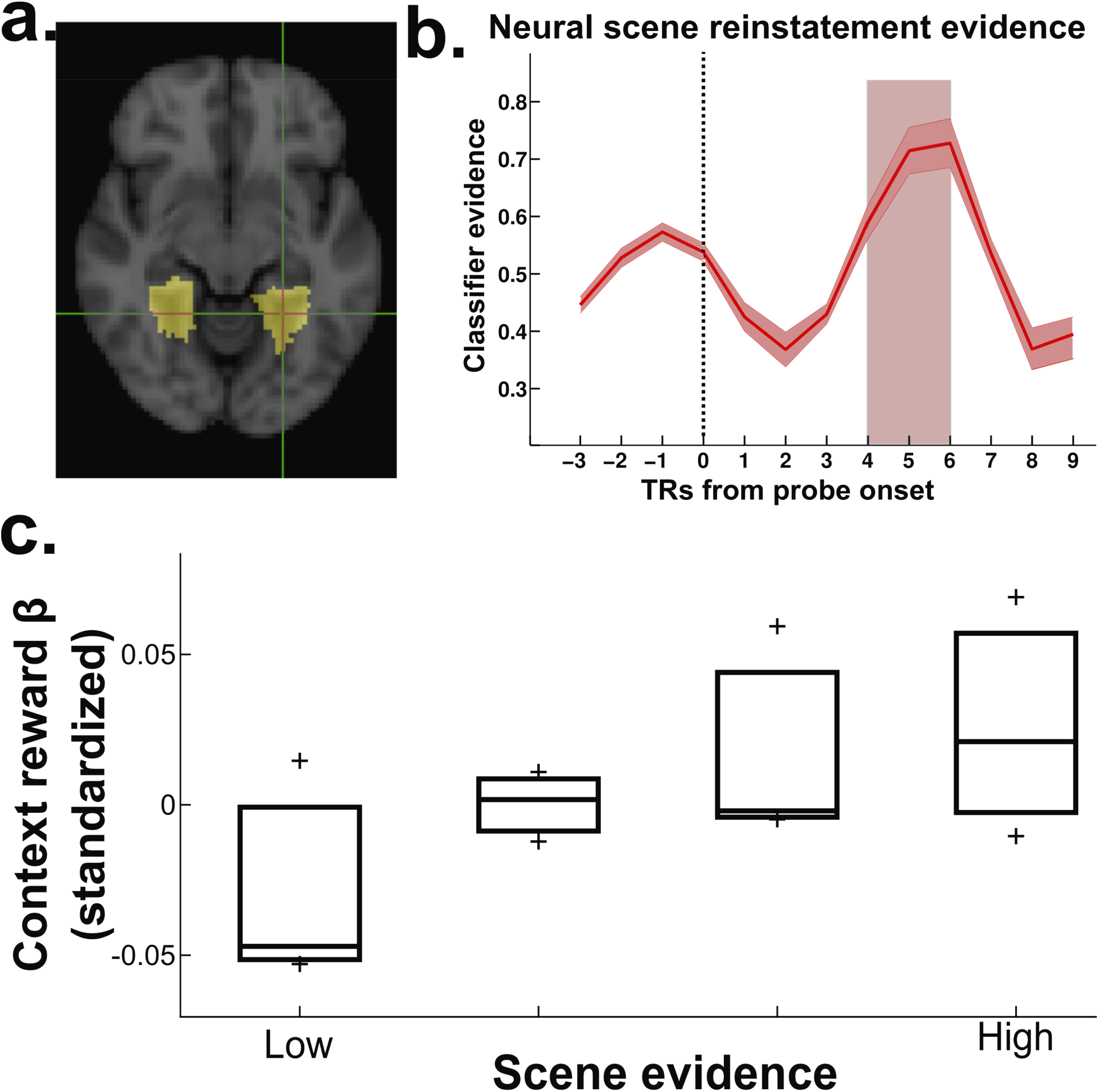
Context reward effect is mediated by scene reinstatement. **a.** We trained a classifier to discriminate between scenes, scrambled scenes, and objects, on the basis of activity in a scene-preferring region of posterior-parahippocampal cortex. **b.** Across subjects and trials, we selected the post-probe timepoints that showed elevated classifier evidence for scenes (.(* P < 0.05) **c.** For each participant, we split choice trials following the probes into quartiles based on scene evidence during the selected timepoints. Repeating the regression of Figure 2 for each quartile of trials, we found that context reward effect was greater when scene evidence was higher (mean slope 0.041, SEM 0.0114, *t*(31)=3.5734, *P*=0.0012; plotted: for each bin, the median, +/- one quartile, across participants, of the standardized regression coefficients for the context reward effect).

We then split probe trials into quartiles for each participant, based on the average level of scene evidence across the timepoints of interest in each trial. In other words, although these timepoints were selected because they showed elevated scene evidence on average, our analysis of interest relied on the variance in scene evidence across probe trials – specifically, we assessed whether scene evidence on probe trials predicted the effect of context reward on choices following those probes. For each quartile, we again ran the behavioral regression above, and calculated the size of the context reward effect, measured as the standardized regression weight applied to the context reward regressor computed as above. This regression weight was normalized to account for the different variance in the variable of interest across bins. We found that, on average, the influence of context reward increased along with classifier evidence for scenes (Figure 3c, difference between lowest and highest quartile significant – *t*(31)=−3.2756, *P*=0.0026; Within subjects, a linear trend across the quartiles was positive and reliable – mean slope 0.041, SEM 0.0114, *t*(31)=3.5734, *P*=0.0012).

#### Context reward is specifically modulated by scene evidence in PPA

To confirm that scene reinstatement specifically modulated context reward, we repeated the analysis in Figure 3c for each of our regressors of interest. We found that scene evidence did not reliably modulate any of the other regressors of interest (all *P* > 0.14; Figure S5). Similar analyses excluded the possibility that our results could be explained by univariate activity or classifier evidence in several control regions (Figures S6 and S7).

#### Reinstatement of individual scenes predicts the influence of context reward

We next examined whether the patterns reinstated at probe trials were specific to individual room contexts. To test this, we first produced, for each participant and each context room, a template pattern consisting of the average PPA activity across all trials within that room. We then computed, at each probe trial that evoked a past decision, the average pattern in PPA across our timepoints of interest (Figure 3b). The evoked patterns were consistently in favor of the reminded room. On high-confidence, correct trials, the average (fisher-transformed) correlation between the pattern at probe and the room-pattern matching the context of the reminded trial was 0.0377 (SEM 0.0047, *t*(31)=7.9494, *P*=5.6502e−09 across subjects), while the average correlation with the non-reminded contexts was 0.0015 (SEM 0.0014, *t*(31)=1.0207, *P*=0.3153). Across subjects, the correlation with the reminded context was reliably larger than that with non-reminded contexts (*t*(31)=6.7397, *P*=1.5248e−07).

We next incorporated this specific-scene measure into our regression, where it improved our estimation of the effect of context reward on choice. First, we turned the correlation values at each trial into a distribution of weights across the six possible contexts, by first adding one to each value (so they were all positive values within the range from zero to two), and then dividing each by the sum across all correlations. This produced a real number, between zero and one, reflecting the relative strength of evidence for reinstatement of each context room. Then, we computed the context reward regressor again for all six contexts, and multiplied these values by the corresponding specific-scene evidence, resulting in a distribution of context reward values scaled by the reinstatement evidence for each context. Finally, the sum of these scaled values was entered into the regression as a reinstatement-weighted version of the original context reward regressor.

Consistent with the idea that the specific content of reinstatements was a strong influence on choices, the regression weight assigned to the evidence-scaled context reward regressor (*β_RCR_*) was reliably positive (mean=0.3851, SEM 0.0863, *t*(31)=4.4611, *P*=9.9968e−05). The regression model containing the new, evidence-scaled, version of the context reward was consistently better than the original model (average difference in R^2^=0.0041, SEM 0.0016, *t*(31)=−2.6794, *P*=0.0117). When both versions of the regressor were placed alongside each other in the same regression model, the original variable was assigned essentially no regression weight (average *β_CR_* =0.0141, SEM 0.0472, *t*(31)=0.8676, *P*=0.3923), while the scene-specific scaled variant remained a strong and consistent predictor of choices (mean *β_RCR_* =0.3923, SEM 0.0877, *t*(31)= 4.4732, *P*=9.6597e−05). Supporting the hypothesis that these reinstatements reflect memory retrieval [2,8], we observed that the entropy of each trial’s distribution of specific-scene reinstatement weights was positively correlated with activity in hippocampus (Figure S8).

Together, these results affirm a specific and measurable role for context-aware episodic sampling during decisions for reward.

## Discussion

Context is a critical aspect of episodic memories. Events do not happen in isolation – the memories of our lived experiences are necessarily situated within a web of associations with internal and external state: where they happened, who they happened with, and what else happened in relation. Items can cue retrieval of contextual features and vice-versa [5]. When we bring our memories of the past to bear in deciding what actions to take in the present, it stands to reason that this rich contextual web would affect which memories are recalled and – through this – what decisions we make. However, previous work – even work investigating the use of episodic memory in decisions – has not examined the impact of contextual associations on choice.

In this study, we investigated how memory for the context of past choice outcomes can affect present decisions for reward. We observed that decisions were biased by incidental memory probes that reminded participants of past choice trials. We observed a separate influence of both reward information on the reminded trial, and of reward information on other trials that shared context with the reminded trial. This influence of reminders is not captured by RL models, and the influence of context in particular is a novel prediction of episodic sampling [2], tested here for the first time. Across studies, the effect of the reminded context on choices was consistently greater than that of the individual reminded trial. This relationship matches our proposed sequential sampling model, where successive samples after that of the reminded trial are drawn from a linked context whose reward statistics were designed to run counter to the probed trials (Figure S3). The effect of both the reminded trial and its associated context were reduced in the second experiment. However, as predicted by the episodic sampling model, the effect of reminded context remained proportionally stronger than the reminded trial, and, critically, a significant influence on choices (Figure S4).

To investigate the neural mechanism that gives rise to this effect, we used pattern classifiers trained on fMRI data to produce a continuous neural measurement of evidence for whether participants reinstated context from memory. Critically, we showed that this neural measurement predicted the size of the behavioral effect: The extent to which participants bring to mind the context of past episodes was correlated with the influence of context reward on decisions. These results are consistent with computational models of temporal context memory [9] – a central prediction of these models is that, when context is reinstated, this leads to to additional memories being recalled from the same context [5]. The present results tie together this effect of context on memory recall, and the neural mechanisms that mediate the effect, with recent findings that support a role for episodic memory recall in deliberative decisions for reward [2, 10].

The mechanism presented here is distinct from, but complementary to, the mechanism thought to underlie previous observations of episodic memory’s involvement in decisions via spreading associations [11]. In that study, Wimmer & Shohamy showed that pairing rewards with unvalenced images can “spread” that reward (during learning) to other unvalenced images that had been associated with the first, and that this second-order reward association can induce a preference for the latter image. Our experiment design has a more complex associative structure that makes this kind of “reward spreading during learning” account highly unlikely. To get spreading during learning, we would need to posit that – when participants make choices late in the context (e.g., choosing the blue deck and getting rewarded) – they mentally activate objects shown early in the context (e.g., binoculars) and associate those objects to the blue deck getting rewarded. This is implausible because the only route for the binoculars to be activated is via their link to the context image, and the binoculars were just one of tens of items that were linked to this context; this high “fan out” of associations from the context image makes it unlikely that participants will strongly activate binoculars or any other single previous item from the context. In any case, even if this did happen infrequently, it is highly implausible that it would happen enough to explain our finding that indirect, contextually mediated associations exert a *stronger* effect on subsequent choice than direct associations. Throughout the paper, we have argued that – rather than “spreading” of associations between stimuli and reward during learning – our results arise from dynamic estimation of that value at the time of choice, based on contextually-mediated sampling. The distinction between these two types of memory-guided decisions reflects the proposed division [12] between “retrospective” (referring to the Wimmer & Shohamy finding) versus “prospective” (this study, also Bornstein et al. [2]) integration. We provide here the first demonstration that such prospective integration is mediated by context-guided sampling from episodic memory, and that this context information induces sequential dependencies in the integration process. This case for prospective, dynamic estimation of value through sampling is convergently supported by our regression modeling (showing that indirect, contextually-mediated effects are larger than direct effects, and that these effects cannot be explained by incremental learning), simulation work (showing that our context-guided sampling model can account for context effects exceeding direct effects; Figures S3 and S4), and neural data (showing that variance in retrieval of specific scene contexts at the time of choice predicts how strongly context memory affects decision behavior).

A key contribution of the present work is that it sharply distinguishes the pattern of choices arising from episodic sampling from those predicted by model-free RL. Our previous work assumed only that episodic memories were drawn according to their recency, yielding a dependence of decisions on past experience that follows the same qualitative form as that of incremental RL [2]. If that recency-dependent sampling was the only way that episodic memories translated to decisions, then it could be argued that model-free RL fits to behavior capture an approximate, average form of the underlying mechanism. However, the discovery that samples depend in part on context undermines the generalizability of that analogy, because context can induce sequential dependencies into the sampling process that are not explainable in terms of simple recency. Our results suggest that, to estimate the influence that a given past trial will have on the current decision, we must know not only its age, but also the relative likelihood that it might be brought to mind by the recall of other past trials. In this study, the context was made visually explicit, but in natural environments, contextual links between episodes may arise from a wide array of external or internal associations [13]; as such, we expect these contextual effects on decision-making to be ubiquitous in everyday life.

More generally, the present findings pose a challenge for economic approaches to modeling decisions. This is because standard economic models eschew consideration of the underlying mechanism, instead focusing exclusively on inferring stable preferences as “revealed” via actual choices [14]. Contrary to this view, our results suggest that choices do not always depend solely on stable preferences – instead, they are constructed dynamically at the time of decision [15, 16], and critically via a process that draws on a complex web of contextual associations that might be only incidentally related to past decisions of the same kind. This idea considerably complicates models of decision making, but it also provides a way forward: By drawing on our understanding of the cognitive and neural mechanisms giving rise to decisions (here, episodic memory retrieval and contextual reinstatement), we account in a principled way for variance in choice behavior that would otherwise be attributed to noise.

## Data and code availability

The data that support the findings of this study are available on reasonable request from the corresponding author (A.M.B). The data are not publicly available because they contain information that could compromise research participant privacy/consent. In the near future, they will be de-identified at the level of contemporary best practices and placed in a public repository, which will be linked to at the below GitHub URL. Standard software packages (SPM8 and FSL 5.0.4) were used for processing the MRI data in addition to custom Matlab scripts. Custom-written analysis code is available upon reasonable request to the corresponding author (A.M.B). Context-aware sampling model code is publicly available at the corresponding author’s GitHub repository (https://github.com/aaronbneuro/neurocode).

## Acknowledgements

The authors wish to thank Jeremy Manning for providing localizer code and stimuli, A. Schapiro, A. Rangel, J. Poppenk, M. deBettencourt, S. Chan and Y. Niv for fruitful discussions, and M. Aly, C. Honey, A. Shenhav and anonymous reviewers for helpful comments on an earlier version of the manuscript. This publication was made possible through the support of a grant from the John Templeton Foundation (Grant ID #57876; K.A.N). The opinions expressed in this publication are those of the authors and do not necessarily reflect the views of the John Templeton Foundation.

## Author Contributions

A.M.B and K.A.N designed the experiment; A.M.B ran the experiment; A.M.B analyzed the data; A.M.B and K.A.N wrote the paper

## Competing Financial Interests Statement

The authors declare that they have no competing financial interests.

## Online Methods

### Participants

23 participants (12 female, mean age 24, range 18-50) performed the task in Experiment 1. Three were excluded for failing memory test criteria (object recognition memory in Phase 2 at d’ < 1, or source recognition performance during Phase 3 not significantly different from chance), leaving 20 participants included in the analyses presented here. 38 participants (21 female, mean age 25, range 18-64) performed the task in Experiment 2. One was excluded for excessive motion during the scan, one was excluded for falling asleep during the scan, and four were excluded for a programming error that caused unrecorded responses, leaving 32 participants included in the analyses presented here. All participants were free of neurological or psychiatric disease, and fully consented to participate. The study protocol was approved by the Institutional Review Board for Human Subjects at Princeton University.

### Task

The experiment was controlled by a script written in Matlab (Mathworks, Natick, MA, USA), using the Psychophysics Toolbox [17]. Participants performed a series of 300 choices between three differently-colored card decks with continuously changing probabilities of reward.

The experiment proceeded in four phases. In *Phase 1*, the *Contexts* phase, 180 choice trials were presented across six consecutive “rooms” of a virtual casino (Figure 1). Rooms were distinguished by the presence of one of six background images of natural scenes. Choices resulted in the top card of the chosen deck being turned over to reveal a trial-unique picture of an everyday object, followed by the presentation of a reward of $10 or $0 (described in detail below, *Choice trials*). The probability that each deck would deliver a $10 reward changed on each trial. This probability was generated according to the procedure described under *Payoffs*. Following Phase 1, participants were asked to rest for as long as they needed, and to indicate their desire to continue by pressing any button twice.

In *Phase 2*, the *Probes* phase, participants performed 120 additional choice trials, along with 60 memory probe trials interspersed at pseudorandom intervals where we tested recognition memory for objects from Phase 1 (see *Phase 2 recognition probes* below). During this phase, participants were told that they had entered a seventh, “unfinished”, room of the casino. In this seventh room, the screen no longer contained a background scene image. Choices in the seventh room did not return object pictures, but continued to be rewarded according to slowly-varying payoff probabilities.

In *Phase 3*, the *Source Recognition* phase, participants answered 50 source recognition memory questions in which they were asked to match a previously-encountered object image to the scene (context) in which it had appeared (in Experiment 1 the choice was out of all six scenes, while in Experiment 2 the choice was among three presented options to fit the constraints of the MRI button box and for clarity of identification on the projected screen).

Lastly, in *Phase 4*, the *Localizer* phase, participants performed a blocked, one-back image repeat detection task. This task was used to identify fMRI responses to three image categories: objects, scenes, and scrambled scenes. The detailed timing and structure of trials in each of these four phases are described below.

Prior to the experiment, participants were given written and verbal instructions as to the types of trials, the payoff probabilities, the button presses required of them, and the rules for determining the final payout. They were told that the decks had different probabilities of paying out, that these probabilities would continually change, and that the probabilities of each deck paying out were independent of each other (as were the outcomes themselves). They were also told that that no aspect of the decks or choice process would change when traveling between rooms – in other words, one could not expect payoff probabilities to shift suddenly when the rooms did – but that the payoff probabilities would occasionally change dramatically, in addition to continuously changing slowly. Instructions emphasized that there was no pattern linking the content of the object pictures to their dollar value or deck. Participants were not told that the Phase 2 memory probe trials should have an effect on their choices, nor was any effect implied. They were, however, told that their final payout would depend in part on their later memory linking the object pictures to the room scenes. To aid their memory, participants practiced, and were encouraged to use, an elaborative encoding strategy in which they would construct, but not vocalize, a sentence describing the object being used in the background scene (e.g. for the object-scene shown pair in Figure 1, an example sentence might be “I use these binoculars to look at the city skyline.”). After participants read the instructions, the experimenter verbally administered a quiz testing their knowledge of the payout rules, room structure, and encoding strategy.

Once in the scanner, participants performed four practice choice trials and one practice memory probe trial, all unscanned, before beginning the main experiment. If participants failed the practice memory probe trial, or expressed a desire to practice again, the practice trials were repeated until both the participant and operator were satisfied.

#### Choice trials

On each choice trial in Phase 1 and Phase 2, participants were presented with three card decks, colored Red, Green, and Blue (order pseudorandomized across participants), arrayed across a green table along the top of the screen (Figure 1a). In Phase 1, the background of the screen contained a picture of one of six outdoor scenes. The background scene remained consistent for 30 consecutive trials, then changed at the onset of next “room”. Participants were given three seconds to make a choice between the decks. Decks were chosen by pressing the “1”, “2”, or “3” key in Experiment 1, and the buttons under the index, middle, or ring finger in Experiment 2, corresponding to the decks from left to right. When a choice was made, the unchosen decks were hidden and the chosen deck was isolated on the green table, and remained so until the end of the three second choice period. For Phase 1 only, the top card of the chosen deck was “turned over” to reveal a trial-unique picture of an everyday object. This picture remained on the screen for two seconds, and then the card was turned back over. The isolated deck remained on the screen and a reward value was displayed – either $10 (a picture of a US $10 bill) or $0 (a phase-scrambled version of the same bill). The reward value remained on the screen for 1.5 seconds, followed by a blank screen for an inter-trial-interval (ITI) of length varying between 0.5 and 8 seconds, mean 1 second, selected from a truncated, discretized exponential distribution generated pseudorandomly for each participant. Between rooms, participants were shown a screen with the name of the next room, and a countdown from four seconds before the next room began.

#### Payoffs

For Phase 1 and Phase 2, choices resulted in rewards with amounts selected according to continually changing probabilities. The probability that each deck I would pay out $10, *π_i,t_*, changed independently on each trial according a decaying Gaussian random walk with reflecting bounds at 5% and 95% (Figure 1b). Specifically, for each deck *i*, payoffs were computed according to Equation 1

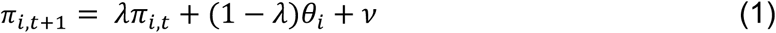

The initial values of the payoff probabilities, *π*_*i*,0_, were set to 60%, 30% and 10%, assigned pseudorandomly without replacement to each deck. The value of the stickiness parameter λ was 0.6, the drift target *θ_i_* was set to the initial payoff for each deck, and the diffusion noise *ν* was zero-mean Gaussian with standard deviation *σ_d_*=8. For the first three trials of each room, the stickiness parameter was temporarily set to 0.95, to ensure that outcomes affirmed to the participants that the preceding payoff probabilities carried through to the new room – in other words, that the decks remained the same, despite changing rooms. Between the tenth and eleventh trial in each room, the targets *θ_i_* of each payoff timeseries were shuffled such that the deck that previously had the highest payout would no longer have the highest payout. Critically, memory probe images were chosen exclusively from the first ten trials of each room. In the seventh room, the payoffs continued drifting as above, with drift targets continuing to swap every thirty trials.

#### Phase 2: recognition probes

In the seventh room (the *Probes* phase) the series of choices was interrupted at pseudorandom intervals by 60 recognition memory probes (Figure 1b). Participants were questioned on their memory for an object photograph. Fifty of the probed photographs had been previously presented on a choice trial during Phase 1; the remaining 10 were novel lures. The probe images were chosen from the first ten trials of the context rooms, because these trials had different payoff values than did the other 20 trials in each room. This feature allowed us to distinguish the influence of memories for the reminded trial from the influence of the reminded context. Participants were instructed to press keys indicating their memory and their confidence level: “1” (indicating highly confident that it was an image they had seen before) through “4” (indicating highly confident that this was an image they had not seen before). For MRI experiment 2, buttons were numbered left to right for the fingers on the right hand, from index finger “1” to pinky finger “4”.

Correct responses – “1” or “2” for previously seen images, or “3” or “4” for images that were not displayed on a previous trial – were rewarded with $0.25 added to the participant’s total payout. This additional reward was indicated by a photograph of a US quarter with a green ‘+’ to the left. Incorrect responses resulted in $0.25 being deducted from the participant’s total payout, indicated by a red ‘-’ to the left of an image of a US quarter. Memory probe rewards were displayed for two seconds.

Rewards for memory probes accumulated over the course of the entire task, rather than for randomly selected rounds – so the total payout could be reduced or increased by as much as $15.00. Probe images remained on the screen for up to three seconds – if no answer was entered in that time, the trial was scored as incorrect.

#### Phase 3: Source recognition test

Before the experiment began, participants were instructed to remember as many of the object pictures as possible, along with their associated rooms. Their memory for these pairings was tested in 50 post-task source recognition probes. Post-task memory probes were drawn from the set of pictures shown during Phase 1 that were also tested in Phase 2 recognition memory probes. Participants were presented with an object picture and candidate rooms (all six in Experiment 1, but only three in Experiment 2 to restrict responses to the one-handed button box used in fMRI), and asked to select the room in which they first saw the object photograph. Each incorrect answer reduced the number of $10 rewards in their pile. The final payout was then determined as the sum of two pseudorandomly selected choice trials, drawn from the set of trials that remained after removing $10 rewards according to the results of the post-task source recognition test.

#### Phase 4: Localizer

To allow us to localize regions of cortex preferentially active during processing of scene images, participants performed a 1-back image repeat detection task. During this localizer task, images were presented in mini-blocks of 10 images. Stimuli in each mini-block were chosen from a large stimulus set of pictures not used in the main experiment, belonging to one of three categories – objects, scenes or phase-scrambled scenes. Images were each presented for 500ms and separated by a 1.3s ISI. Eight of the images in each block were trial-unique, and two were repeats. Repeats were inserted pseudorandomly, according to a uniform distribution. A total of 30 mini-blocks were presented (10 per each category), with each mini-block separated by a 12 second inter-block interval.

### Imaging methods

Data were acquired using a 3T Siemens Skyra scanner with a 20-channel volume head coil. We collected two functional runs with a T2*-weighted gradient-echo echo-planar sequence (37 oblique axial slices, 3mm isotropic resolution, echo time 27.0 ms; TR 2080 ms; flip angle 64; field of view 192 mm). The first four volumes of each functional run (8.32s) were discarded to allow for T1 equilibration effects. We also collected a high-resolution 3D T1-weighted MPRAGE sequence for registration across participants to standard space. Functional image preprocessing was performed using FSL (FMRIB Software Library version 5.0.4; [18]). Anatomical images were coregistered to the standard MNI152 template image, then individual participant functional images were coregistered to the realigned anatomical images. The transformation matrices generated during this coregistration process were used to transform Region of Interest (ROI) images (described below, ROI definition). Functional images were motion corrected and spatially smoothed using a 5mm full-width half-maximum Gaussian kernel prior to analysis. Data were scaled to their global mean intensity and high-pass filtered with a cutoff period of 128s. Pattern analyses were performed using the Princeton Multi-Voxel Pattern Analysis Toolbox (MVPA Toolbox; http://www.pni.princeton.edu/mvpa) and custom code implemented in MATLAB.

### Behavioral analysis

#### Regression analysis

To examine the influence of past trials on choice in Phase 2, we conducted a regression analysis relating the outcomes of past choices to the choice made on the current trial. This regression included outcomes both from choice trials where rewards were directly experienced, and from trials evoked by memory probes.

We constructed the following design matrix three times, once with each deck – red, green, and blue – as the given deck of interest. We first entered into the regression the *identity* of the deck chosen on the previous trial: 1 for the given deck, 0 for others. Next, we entered variables describing the directly received rewards. If a reward was received after choosing the given deck on trial *t* − *τ*, this was coded as a 1 in regressor *τ*, element *t*. If no reward was received after choosing the given deck, this was coded as a 0.

Next, we included two variables coding aspects of the reminded trial (i.e., the trial cued by the memory probe, if there was a memory probe on the preceding trial). The first regressor coded the evoked identity of the deck chosen on the reminded trial (again, 1 for the given deck, 0 for others), and the following regressor coded for the evoked reward received on the reminded trial.

#### Context reward

The final regressor coded for the *context reward*, the net reward actually experienced for choosing the given deck within the evoked context. As discussed in the *Results*, we designed this regressor to capture the average effect of sampling memories from the probed context.

For each deck, this value was calculated as the number of trials on which the option was chosen and rewarded, minus the number of trials on which the option was chosen and not rewarded, divided by the total number of trials on which the option was chosen. Explicitly, for deck *i* in context *C*, this value is:

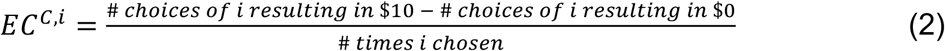

If a deck was not chosen in the evoked context, *EC^C,i^* was set to zero. If the preceding trial was not a memory probe, the evoked identity, evoked reward, and were all set to zero. In the dependent variable, choices were coded as 1 if the given deck was chosen and 0 otherwise.

The regression was thus in the following form:

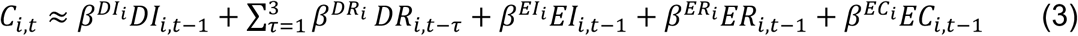

where *C*_*i,t*_ specifies whether deck *i* was chosen at trial *t*, *DI* is ‘directly experienced identity’, *DR* is ‘directly experienced reward’, *EI* is ‘evoked identity’, and *ER* is ‘evoked reward’ for each trial preceding the current choice, and also for the given deck *i*. Following the patterns observed in our previous study [2], effects of memory and the identity of the previous deck are specified for the previous trial, while direct reward receipt is specified for the preceding three trials.

In total, there were eight columns in the design matrix – the seven predictor variables just described, plus the constant term, and 360 rows – one for each of the 120 Test-phase choices, each specified three times coded for the three decks of interest. The resulting regression weights – indicating the degree to which the current choice was influenced by choices and rewards on a given evoked or directly experienced trial or context – were treated as random effects and tested against zero across the population by two-tailed *t*-test. See Table 1 for a concrete illustration of how the design matrix was constructed.

**Table 1:** Example regression design matrix. This table depicts the rows of the design matrix that code for the fourth trial, given the following scenario: Red was selected on the three preceding choice trials, the first two of which resulted in rewards; after the third Red trial, a memory probe was presented that evoked a trial on which Blue was chosen, in a context in which Blue was chosen on 12 trials and rewarded on 7, and Green was chosen on 18 trials and rewarded on 13. The regression design contained three rows for each trial, each reflecting the contribution of the independent variables to the probability of picking a given deck (Red, Green, or Blue). The independent variables code for the presence of choice-relevant information on recent trials. For instance, the first column (*DI_-1_*) indicates whether the deck of interest was selected on the most recent choice trial. The second column (*DR_-1_*) indicates whether the deck of interest was associated with reward on the most recent choice trial. The third and fourth (*DR_-2_, DR_-3_*) columns indicate whether the deck of interest was associated with reward on the preceding two choice trials. If the most recent trial before the current choice was a memory probe, the fifth through seventh columns contain indicators of the choice and value information on the trial evoked by the probed image: respectively, the identity of the deck chosen on the reminded trial (*EI_-1_*), whether or not that choice resulted in reward (*ER_-1_*), and the reward received for choosing the given deck across the room in which the reminded trial took place (*EC_-1_*) (Equation 2). The dependent variable predicted by the regression was a 1 or 0 coding for whether the deck of interest was selected on the current trial (trial 4 in this example). The resulting regression coefficients reflect the contribution of each variable to the probability of choosing as the participant did.

| Deck, Trial | $DI_{-1}$ | $DR_{-1}$ | $DR_{-2}$ | $DR_{-3}$ | $EI_{-1}$ | $ER_{-1}$ | $EC_{-1}$ |
| --- | --- | --- | --- | --- | --- | --- | --- |
| Red, 4 | 1 | 0 | 1 | 1 | 0 | 0 | 0 |
| Blue, 4 | 0 | 0 | 0 | 0 | 1 | 1 | 0.17 |
| Green, 4 | 0 | 0 | 0 | 0 | 0 | 0 | 0.44 |

#### Reinstatement-scaled context reward

We also ran the regression analysis using a version of the context reward regressor that was augmented with neuroimaging evidence for reinstatement of each specific scene context (for details on the neuroimaging evidence calculation, see *Specific-scene patterns*, below).

In this analysis, the *EC* regressor (Equation 2) was recomputed six times at each choice following a memory probe, once for each of the context rooms, producing *EC^C,k^*. In other words, the regressor was computed as though the probe had reminded the participant of each room.

We next incorporated the scene-specific reinstatement measure into our regression. First, we turned the correlation values at each trial into a distribution of weights across the six possible contexts. We did this by adding 1 (to account for negative correlations) to the correlation values, and then dividing the resulting values by the sum across all correlations. This resulted in a real number, between zero and one, indicating the relative degree of match between activity at probe and the template for each context.

The six context reward values, *EC^C,k^*, were each multiplied by our estimated probability that the participant was reinstating the given context *C*^*k*^. The sum of the six probability-weighted context rewards was then entered into the regression in place of (or, in the second analysis, alongside) the original *EC* regressor.

#### Context-aware sampling model

To investigate how single-trial and context reward trade off with each other as the number of past episodes sampled increases, we simulated the task as performed by a context-aware episodic sampling model. In this simulation, all choices are made using episodic sampling alone (no influence of model-free values), to clearly isolate the influence of changing the number of samples. In episodic sampling, option values are estimated using the values encountered on one or more past episodes, with the likelihood of sampling a given episode diminishing exponentially with its recency. The context-aware episodic sampling model augments this idea, by positing that additional samples after the first are (with some probability) selected uniformly from the same context as the preceding sample.

The model maintains a cache of episodes representing each experienced trial. When subjects respond correctly to a valid memory probes, the model with some probability “reinstates” the episode by copying the reminded trial to the front of the cache–thus, making it more likely to be drawn when evaluating options during the next choice. If the subject’s correct response to the memory probe is of high confidence, then the context of the probed trial is included in the episode copied to the front of the cache.

We used this model to simulate subjects who used different numbers of samples from episodic memory to make decisions. The model had four parameters which were fixed across all simulations: *α_direct_*, or the decay rate on temporal recency; *α_evoked_*, or the probability of reinstating evoked trials because of memory probes; *β*, the softmax temperature; *β_p_*, the choice perseveration term; and *π*, the likelihood of drawing sample *k* from the same context as sample *k*-1 (as opposed to based on temporal recency).

A final parameter, the number of samples drawn, was varied between 1 and 15. For each fixed number of samples, we simulated 1,600 subjects (50 groups of 32) each performing an instantiation of the task. The simulated subject’s parameters *α_direct_*, *β*, and *β_p_*, were set to those fit to the real subjects with the number of samples equal to 1, and *π* was set to 1.

For simplicity, in the first simulation *α_evoked_* was set to 1. This assumption was relaxed for our second set of simulations, during which we varied *α_evoked_* between 0 and 1 to illustrate the impact of changing this parameter. The simulated subjects were programmed to make, on average, the same proportion of low confidence and incorrect responses as did the subjects from Experiments 1 and 2.

#### Incremental learning models

To compare our context reward model to the others, we generated the timeseries of reinstated context reward values that would be learned according to three different specifications of model-free RL.

The first variant followed the traditional method, learning the value of each card deck without regard to changes in room context. Specifically, at each step, the value for the chosen card deck, *Q_B_*, was updated according to the reward received on that trial, *R_t_*, and the learning rate *α*:

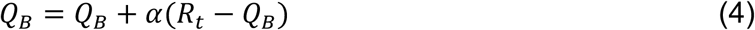

We refer to this as the *standard* model. The second variant reset the value of each card deck when the room changed, giving separate values *Q_B,C_* for each room-context, following Equation 5.

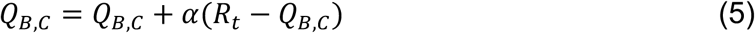

We refer to this as the *room-reset* model. A third variant also reset action values, but this time it did so at a variable trial number within each room (e.g. 1, 2, 3, or more trials after room change). We designed this model to account for the possibility that participants took note of payoff reversals and discounted information prior to the switch [22]. We refer to this model as the *reversal* model.

The three incremental learning models were fit to choices in rooms one through six, using maximum likelihood estimation of parameters. To compare the model likelihoods, we first transformed them using the Bayesian Information Criterion (BIC; [23]), which penalized the third model for its additional parameter.

The best-fitting model was used to generate a replacement for the context-reward regressor as specified in *Regression analysis*. The resulting regression model weights were compared to the original (Figure 2).

### Imaging analysis

To identify neural markers of context reinstatement, we first defined, using MVPA [19], the pattern of BOLD activity in posterior parahippocampal cortex that indicated participants were processing “scene” images. We then looked for evidence that this pattern was reinstated following probe trials. We reasoned that greater evidence of scene reinstatement would indicate that participants were recalling the context of the probed image (note that no scene images were present during Phase 2), and thus would show an increased influence of other trials from this context on their decision-making.

ROI definition. We identified a region of interest consisting of voxels that (across the group) showed preferential activation to scene images, using the following procedure. First, for each participant, we performed a GLM analysis of the localizer phase data, and identified voxels selectively responding to scenes versus other categories (univariate contrast, scenes > scrambled_scenes|objects). For each participant, we selected clusters in the posterior parahippocampal region (matching the reported Parahippocampal Place Area (PPA); [20]) that were significant at *P*<0.005, uncorrected. Next, each per-participant voxel mask was binarized; all above-threshold voxels were set to 1. This mask was warped to match the group average anatomical so that each participant’s mask could be aligned and the collection averaged. The resulting group-space masks were added together and the summed image thresholded to include all voxels present in more than 90% of participants. This final group ROI was then warped back to the individual participant space, and the result used as a mask for pattern classifier analyses.

To permit various control analyses, we followed a parallel procedure to identify a region of interest that preferentially responded to the scrambled scenes used in our localizer task. For this ROI, rather than selecting clusters within an anatomical area of interest, we simply used the contrast mask from across the entire brain.

#### Category-level pattern classification

We trained a classifier to identify patterns of activity indicative of participants processing pictures of scenes. We first extracted, across the localizer task, activity of all of the voxels in the above-defined scene-responsive ROI. These labeled data were used to train an L2-regularized multinomial logistic regression classifier to predict scene versus scrambled scene labels. The regularization parameter was set to 0.1, but the results were insensitive to varying this parameter by several orders of magnitude in either direction.

The trained classifier was then applied to activity after each probe trial. For each TR of interest, at each probe trial, the classifier provided a measure of the probability that participants were processing scenes; we refer to this real-valued number as scene evidence. We first selected as TRs of interest those timepoints after each probe that reflected peak selectivity to scenes. Because no scenes were on the screen during or after the probes, we treated elevated scene evidence as indicating that participants recollecting contextual information (background scenes) from Phase 1. We compared scene evidence in different conditions and at different time points using paired-sample t-tests; all tests were two-tailed.

Our final analysis involves splitting probe trials into four bins by the amount of classifier evidence for scenes on our selected timepoints of interest. These bins may have different variance within them, which could potentially confound the subsequent regression analysis we perform using these evidence quantities. Therefore, to evaluate the relative contribution of classifier evidence in different quartiles to explaining the context reward effect, we report standardized regression coefficients (Schroeder et al. [21]; Equation 6) that scale the regression weights by the relative variance of the evidence timeseries in quartile *i* as a proportion of the variance of the context reward in that quartile:

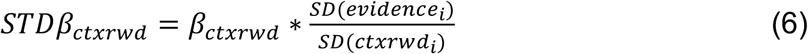

#### Specific-scene patterns

We evaluated whether memory probes caused participants to reinstate the specific context, rather than just a general measure of scene reinstatement, and whether measurements of the contexts that were actually reinstated could improve our predictions of behavior. For each participant, and for each context room, we produced a template pattern of PPA activity in that room, by averaging activity in each PPA voxel across all 30 choice trials. We then computed, at each probe trial that evoked a past decision, the average pattern in PPA across our timepoints of interest (Figure 3b). Next, we computed the fisher-transformed correlations between these trial patterns and each template. These values were then entered into the multiple regression analysis, as described in the section Reinstatement-scaled context reward, above.

## Statistics

A within-subject design was used. Thus, experimental group randomization or blinding was not applicable. To examine the effect of recent and reminded rewards on choices, and parameters across model fits, we performed a multiple regression separately for each subject, and tested the resulting population of beta weights against zero, using one-sample *t*-tests. We compared beta weights within experiments using paired two-sample *t-*tests. We used unpaired, two-sample *t*-tests to compare beta weights across experiments. All *t*-tests were two-tailed, except for the localizer contrast, which used the standard SPM one-sample tests. Data distribution was assumed to be normal, but this was not formally tested before analysis. No statistical methods were used to pre-determine sample sizes. Sample size (20) in Experiment 1 was based on a previous experiment using a similar task [2]. For Experiment 2, sample size (32) was based on our current standard lab practice for fMRI sample stopping criterion, as well as referenced to the sample size in a previous fMRI study performed in our lab (14) that also used classifier evidence of scenes as a signature of context reinstatement [7], which here was more than doubled to account for the finer degree of distinction required for these analyses. Incremental learning models were compared on the basis of their likelihoods (summed log choice probabilities), using BIC [23] to penalize the third model for its additional parameter.

## References

[1] Brian Lau and Paul W Glimcher. Dynamic Response-by-Response Models of Matching Behavior in Rhesus Monkeys. Journal of the Experimental Analysis of Behavior, 84(3):555–579, nov 2005. ISSN 0022-5002. doi: 10.1901/jeab.2005.110-04. URL http://www.pubmedcentral.gov/articlerender.fcgi?artid=1389781.

[2] Aaron M Bornstein, Mel W Khaw, Daphna Shohamy, and Nathaniel D. Daw. What’s past is present: Reminders of past choices bias decisions for reward in humans. Nature Communications, 2017.

[3] N D Daw, J P O’Doherty, P Dayan, B Seymour, and R J Dolan. Cortical substrates for exploratory decisions in humans. Nature, 441:876–879, 2006.

[4] Sean M. Polyn, Kenneth A. Norman, and Michael J. Kahana. A context maintenance and retrieval model of organizational processes in free recall. Psychological Review, 116(1):129–156, 2009. ISSN 1939-1471. doi: 10.1037/a0014420. URL http://doi.apa.org/getdoi.cfm?doi=10.1037/a0014420.

[5] Jeremy R Manning, Michael J Kahana, and Kenneth A Norman. The role of context in episodic memory. In M Gazzaniga, editor, The Cognitive Neurosciences, pages 557–566. MIT Press, Cambridge, MA, 2014.

[6] Per B Sederberg, Samuel J Gershman, Sean M Polyn, and Kenneth A Norman. Human memory reconsolidation can be explained using the Temporal Context Model. Psychonomic Bulletin & Review, 18:455–469, 2011.

[7] Samuel J Gershman, Anna C Schapiro, Almut Hupbach, and Kenneth A Norman. Neural context reinstatement predicts memory misattribution. The Journal of Neuroscience, 33(20):8590–5, May 2013. ISSN 1529-2401. doi: 10.1523/JNEUROSCI.0096-13.2013. URL http://www.ncbi.nlm.nih.gov/pubmed/23678104.

[8] Michael N Shadlen and Daphna Shohamy. Decision Making and Sequential Sampling from Memory. Neuron, 90(5):927–939, 2016 doi: 10.1016/j.neuron.2016.04.036. ISSN 927-939 URL http://dx.doi.org/10.1016/j.neuron.2016.04.036.

[9] Marc W. Howard and Michael J. Kahana. A Distributed Representation of Temporal Context. Journal of Mathematical Psychology, 46(3):269– 299, jun 2002. ISSN 00222496. doi: 10.1006/jmps.2001.1388. URL http://linkinghub.elsevier.com/retrieve/pii/S0022249601913884.

[10] Aaron M. Bornstein and Nathaniel D. Daw. Cortical and Hippocampal Correlates of Deliberation During Model-Based Decisions for Rewards in Humans. PLoS Computational Biology, 9(12):e1003387, dec 2013. ISSN 1553-7358. doi: 10.1371/journal.pcbi.1003387. URL http://dx.plos.org/10.1371/journal.pcbi.1003387.

[11] G.E. Wimmer and D. Shohamy. Preference by association: How memory mechanisms in the hippocampus bias decisions. Science, 338:270–3, 2012.

[12] Daphna Shohamy and Nathaniel D. Daw. Integrating memories to guide decisions. Current Opinion in Behavioral Sciences, 5(October):85–90, 2015.

[13] Samuel J Gershman and Yael Niv. Learning latent structure: carving nature at its joints. Current Opinion in Neurobiology, 20(2):251–6, apr 2010. ISSN 1873-6882. doi: 10.1016/j.conb.2010.02.008.

[14] B Douglas Bernheim. On the potential of Neuroeconomics: A critical (but hopeful) appraisal. American Economic Journal: Microeconomics, 1(2):1–41, 2009. ISSN 1098-6596. doi: 10.1017/CBO9781107415324.004.

[15] Elke U Weber and Eric J Johnson. Constructing Preferences from Memory. In S Lichtenstein and P Slovic, editors, The Construction of Preference, pages 397–410. Cambridge University Press, New York, NY, 2006.

[16] Ido Erev, Eyal Ert, and Eldad Yechiam. Loss Aversion, Diminishing Sensitivity, and the Effect of Experience on Repeated Decisions. Journal of Behavioral Decision Making, 21 (May):575–597, 2008. doi: 10.1002/bdm.

## Methods-only References

[17] David H. Brainard. The Psychophysics Toolbox. Spatial Vision, 10(4):433–6, Jan 1997. ISSN 0169-1015. URL http://www.ncbi.nlm.nih.gov/pubmed/9176952.

[18] Stephen M. Smith, Mark Jenkinson, Mark W. Woolrich, Christian F. Beckmann, Timothy E J Behrens, Heidi Johansen-Berg, Peter R. Bannister, Marilena De Luca, Ivana Drobnjak, David E. Flitney, Rami K. Niazy, James Saunders, John Vickers, Yongyue Zhang, Nicola De Stefano, J. Michael Brady, and Paul M. Matthews. Advances in functional and structural MR image analysis and implementation as FSL. NeuroImage, 23 (SUPPL. 1):208–219, 2004. ISSN 10538119. doi: 10.1016/j.neuroimage.2004.07.051.

[19] Kenneth A. Norman, Sean M. Polyn, Greg J. Detre, and James V. Haxby. Beyond mind-reading: multi-voxel pattern analysis of fMRI data. Trends in Cognitive Sciences, 10(9):424–430, 2006. ISSN 13646613. doi: 10.1016/j.tics.2006.07.005.

[20] R Epstein and N Kanwisher. A cortical representation of the local visual environment. Nature, 392(6676):598–601, Apr 1998. ISSN 0028-0836. doi: 10.1038/33402. URL http://www.ncbi.nlm.nih.gov/pubmed/9560155.

[21] L. D. Schroeder, D. L. Sjoquist, and P. E. Stephan. Understanding regression analysis: An introductory guide. Sage, Beverly Hills, CA, 1986.

[22] Timothy E. J. Behrens, Mark W. Woolrich, Mark E. Walton, and Matthew F S Rushworth. Learning the value of information in an uncertain world. Nature Neuroscience, 10(9):1214–21, Sep 2007. ISSN 1097-6256. doi: 10.1038/nn1954. URL http://www.ncbi.nlm.nih.gov/pubmed/17676057.

[23] Gideon Schwarz. Estimating the Dimension of a Model. Annals of Statistics, 6(2):461–464, 1978.

